# Benchmarker: an unbiased, association-data-driven strategy to evaluate gene prioritization algorithms

**DOI:** 10.1101/497602

**Authors:** Rebecca S. Fine, Tune H. Pers, Tiffany Amariuta, Soumya Raychaudhuri, Joel N. Hirschhorn

## Abstract

Genome-wide association studies (GWAS) are valuable for understanding human biology, but associated loci typically contain multiple associated variants and genes. Thus, algorithms that prioritize likely causal genes and variants for a given phenotype can provide biological interpretations of association data. However, a critical, currently missing capability is to objectively compare performance of such algorithms. Typical comparisons rely on “gold standard” genes harboring causal coding variants, but such gold standards may be biased and incomplete. To address this issue, we developed Benchmarker, an unbiased, data-driven benchmarking method that compares performance of prioritization strategies to each other (and to random chance) by leave-one-chromosome-out crossvalidation with stratified linkage disequilibrium (LD) score regression. We first applied Benchmarker to twenty well-powered GWAS and compared gene prioritization based on strategies employing three different data sources, including annotated gene sets and gene expression. No individual strategy clearly outperformed the others, but genes prioritized by multiple strategies had higher per-SNP heritability than those prioritized by one strategy only. We also compared two gene prioritization methods, DEPICT and MAGMA; genes prioritized by both methods strongly outperformed genes prioritized by only one. Our results suggest that combining data sources and algorithms should pinpoint higher quality genes for follow-up. Benchmarker provides an unbiased approach to evaluate any method that provides genome-wide prioritization of gene sets, genes, or variants, and can determine the best such method for any particular GWAS. Our method addresses an important unmet need for rigorous tool assessment and can assist in mapping genetic associations to causal function.

## Introduction

Genome-wide association studies (GWAS) have successfully identified thousands of loci genetically associated with a wide range of human diseases and traits^1^. However, determining the causal variants and genes within these loci remains challenging: the true identities of the causal variants are often obfuscated by linkage disequilibrium (LD) between neighboring variants, and assigning noncoding variants to the genes they regulate has proven difficult. To address these issues, numerous types of algorithms to prioritize the most likely causal variants and genes have been developed^2–8^. Many of these algorithms are based on a simple intuition: when all potentially causal genes or variants are pooled, we expect that those that are actually causal should share more genomic features and/or annotations in common with other causal genes and variants than with non-causal genes and variants. In other words, genomic features or annotations that are shared more strongly than expected by chance in the pool of potential causal genes can be used to prioritize genes and variants. For example, algorithms have been developed that prioritize genes or variants that share similar profiles within gene sets^9^, protein-protein interaction (PPI) or co-expression networks^6,7,10–13^, PubMed abstracts^14^, and regulatory features such as DNase hypersensitivity sites^15^.

Gene prioritization is a critical step in translating genetic discoveries into biological insights, so many methods for gene prioritization have been developed. However, it is not straightforward to compare, or “benchmark,” the performance of these methods and assess which of them produces the most accurate results. Most published prioritization algorithms contain a validation component, but each study takes its own approach to do this, making comparison between algorithms difficult. A common benchmarking approach is to use “gold standard” genes (i.e. genes with a known link to the trait of interest)^9,16^ to calculate a receiver operating characteristic or similar metric. Unfortunately, this strategy relies heavily on prior knowledge of disease etiology and is biased toward well-studied genes in well-characterized biological pathways. In fact, using gold standard genes may actually penalize a method that successfully discovers novel biology (and of course the accuracy of this strategy will suffer if any of the genes classified as “gold standards” turn out not to be truly causal). Another common approach is prospective validation, using a newer (and generally better-powered) data set to benchmark the prioritization results from an older one. One study conducted a large benchmarking effort with this strategy by collecting 42 novel trait-gene associations over six months to use for benchmarking purposes^17^. This methodology, however, requires the existence of multiple independent well-powered GWAS studies, which may not always be available. Another study used Gene Ontology (GO) annotations^18^ and the FunCoup network^19^ to benchmark several network-based prioritization strategies using cross-validation^20^. However, this strategy assumes that a method’s ability to use PPI network connectivity to recover withheld members of a GO gene set is equivalent to its ability to use that connectivity to prioritize causal genes from a GWAS, which may not be the case. Causal genes for a trait likely relate to one another in ways more complex than membership in one gene set, so this analysis may measure the relationship between GO and the FunCoup network rather than the effectiveness of the prioritization methods *per se*.

An ideal strategy would combine the best features of the previously described large-scale benchmarking efforts: (1) cross-validation and (2) the use of GWAS data (rather than external sources of information that can be biased or in many cases nonexistent) as a benchmark. To that end, we propose a novel “leave-one-chromosome-out” strategy for benchmarking, in which the full set of GWAS data is used for both prediction and validation. Specifically, to benchmark one or more methods, we use those methods to prioritize genes on each chromosome in turn, using GWAS data for all the other chromosomes. Next, for each method being benchmarked, we assemble all of the prioritized genes on each chromosome into a single group. Finally, we apply stratified LD score regression^21^ to each group of prioritized genes to determine whether the prioritized group significantly contributes to trait heritability. To compare multiple methods within a given trait, we compare the contribution to trait heritability by genes prioritized by each method. In this way, the GWAS data itself serves as its own control, without the need for incorporating additional data sources, and the use of the leave-one-chromosome-out approach prevents overfitting because association signals are not correlated across chromosomes. This strategy is highly generalizable because it can be applied to any method that prioritizes genes or variants based on their similarity to each other with respect to some feature(s) of interest (e.g. similar patterns of gene set membership, similar epigenetic marks).

## Materials and Methods

### Overview

We first describe the GWAS data we have used to test our method. Then, we describe our approach, which we refer to as Benchmarker; we include an overview of stratified LD score regression. Finally, we discuss the specific prioritization approaches tested.

### GWAS data

We obtained GWAS summary statistics from publicly available resources (Supplemental Table 1). Specifically, we used published summary statistics for height^22^, schizophrenia^23^, inflammatory bowel disease (IBD)^24^, and several lipid measures (low-density lipoprotein [LDL] cholesterol, high-density lipoprotein [HDL] cholesterol, triglyceride level, and total cholesterol)^25^. We also used published UK Biobank summary statistics for body-mass index (BMI), waist-hip ratio adjusted for BMI (WHRadjBMI), skin pigment, red blood cell count, white blood cell count, diastolic and systolic blood pressure, years of education, smoking status, diagnosis of allergy or eczema, age of menarche, and age of menopause^26^. The UK Biobank GWAS each consist of an average of 448,690 European samples analyzed with BOLT-LMM (with the exception of menarche and menopause, which comprise 242,278 and 143,025 female-only samples, respectively).

### Method evaluation

To evaluate a given prioritization method, we assume that variants near and within the set of truly causal genes will be, on average, enriched for heritability. We first apply Benchmarker to a GWAS from which one chromosome has been removed (Figure 1). We consider the top-ranked 10% of genes from the withheld chromosome to be “prioritized.” We repeat this for all 22 autosomal chromosomes and combine all prioritized genes together, which represents the set to be tested.

**Figure 1:**
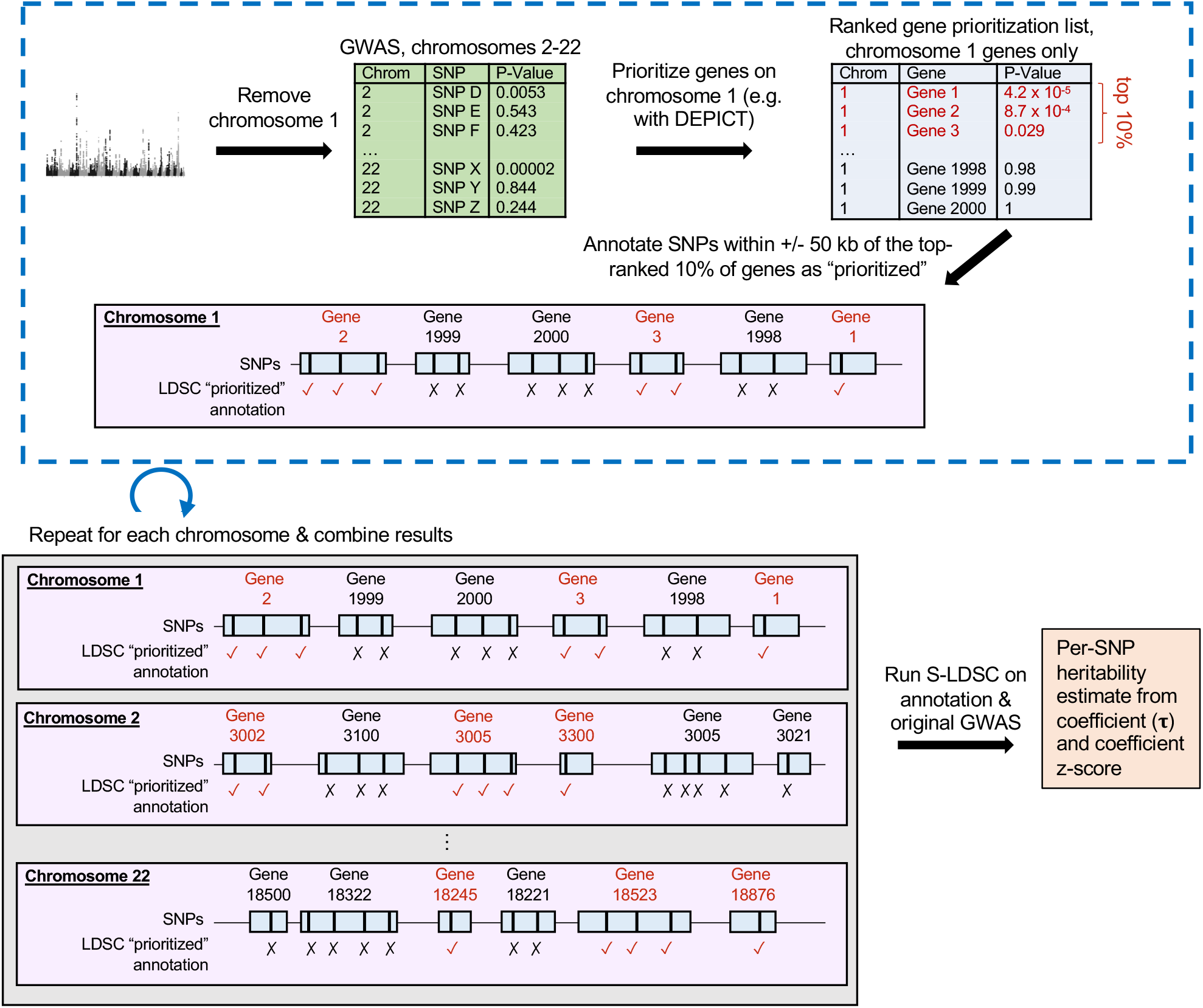
Schematic of the Benchmaker strategy.

To evaluate method performance, we turned to a well-established method: stratified LD score regression^21^. LD score regression is based on the intuition that SNPs with more LD to other SNPs are more likely to tag a truly causal variant (and therefore to have a higher χ^2^ statistic)^27^. An “LD score” for each SNP is calculated by summing its squared Pearson correlation coefficient (r^2^) with all nearby SNPs. Specifically, the LD score of index SNP *k* is given by:

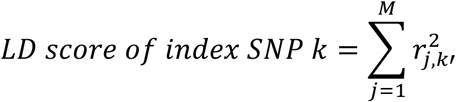

where *M* is the number of SNPs within 1 cM of SNP *k*. The slope of the relationship between LD scores and observed chi-square values, computed in a weighted regression, provides a reliable estimate of overall heritability for a given trait^21^.

Stratified LD score regression (S-LDSC) is an extension of LD score regression that allows for the estimation of the heritability explained by a particular genomic annotation (e.g. coding SNPs, SNPs predicted to be enhancers)^21^. In stratified LD score regression, LD scores are calculated as described above, but only consider the LD of the index SNP with other SNPs in the category of interest *C* (rather than across all SNPs). In our case, *C* represents the set of prioritized SNPs (or SNPs in prioritized genes). For robust and more accurate estimation of the heritability captured by an annotation of interest, it is recommended that a set of annotations of known genomic importance be included in the S-LDSC regression model as conditional covariates. These 53 such annotations are referred to as the “baseline model”^21^ and include annotations such as sequence characteristics (e.g. exon, intron) and cell-type-nonspecific regulatory marks, such as histone modifications. We therefore included the baseline model in each iteration of S-LDSC, as well as an additional category for SNPs that lie within 50 kb of any gene in our prioritization method as a control. Our reference SNPs for European LD score estimation were the set of 9,997,231 SNPs with a minor allele count ≥ 5 from 489 unrelated European individuals in Phase 3 of 1000 Genomes^28^. Heritability was partitioned for the set of 5,961,159 SNPs with MAF ≥ 0.05; regression coefficient estimation was performed with 1,217,312 HapMap3 SNPs (SNPs in HapMap3 are used because they are generally well-imputed).

To evaluate model performance, we focused on two metrics: the regression coefficient τ and its p-value (derived from a block jackknife) for our annotation. τ measures the average per-SNP contribution of the annotation to heritability after accounting for the other categories in the model. We note that, since we used the top 10% of genes for each method, the number of prioritized SNPs for each trait will vary mainly by the average gene length of prioritized genes. To make τ comparable across traits, we normalized by the average per-SNP heritability for each trait (i.e. we divided each estimate by the total trait heritability / the number of SNPs used to compute total trait heritability); we refer to this as “normalized τ”. In previous work^29^, normalized τ may be multiplied by the standard deviation of the annotation; this quantity, τ*, measures the increase in per-SNP heritability per standard deviation increase in the annotation value. For a binary annotation, the standard deviation corresponds directly to the proportion of SNPs assigned to that annotation. For our purposes, we were less interested in the change in heritability associated with a one-standard-deviation increase in the annotation value (which, for a binary annotation, does not have an intuitive interpretation) and more interested in the change in heritability associated with the value of the annotation increasing from 0 to 1, which is captured by normalized τ rather than τ*. We therefore compute normalized τ as an evaluation metric throughout our analyses (this normalization has also been used in Finucane et al. 2018^30^).

To compute p-values for pairwise comparisons, we used a block jackknife to compute the standard error around the difference between two τ estimates within a trait (i.e. τ_A_ - τ_B_). We also calculated meta-analysis p-values from the normalized τ values to determine whether one annotation generally outperformed another across multiple traits. To do this, we selected only one GWAS from groups of obviously overlapping sets of traits; specifically, from the lipid trait group we retained LDL cholesterol and excluded HDL cholesterol, triglycerides, and total cholesterol, and from the blood pressure traits we included systolic blood pressure and excluded diastolic blood pressure.

### Prioritization methods

We began by evaluating DEPICT (release 194)^9^, a method for gene prioritization, gene set enrichment analysis, and tissue enrichment analysis. DEPICT’s primary innovation is the use of “reconstituted” gene sets, which consist of 14,462 gene sets downloaded from multiple databases that have been extended based on 77,840 publicly available expression microarrays^31^. The reconstituted gene sets contain z-scores for each gene in the genome for each of the 14,462 gene sets, representing how strongly each gene is predicted to be a member of each gene set. DEPICT’s gene prioritization algorithm involves 1) identifying all genes in trait-associated loci (referred to as *S*), which in DEPICT are defined as all genes overlapping any SNP with r^2^ > .5 to the index variant and 2) for each of the genes identified in S, assessing its correlation with the rest of the genes in S across the reconstituted gene sets. The stronger the overall correlation a gene has with the rest of the genes in S, the more highly it will be prioritized. We adapted this method for Benchmarker by forcing the prioritization to be genome-wide rather than across only the genes in S (that is, each gene in the genome is compared to the genes in S across the reconstituted gene sets). DEPICT requires a GWAS p-value threshold to define “trait-associated loci.” We used p<10^-5^ for our analyses here, except for a few GWAS for which this threshold caused DEPICT to exceed its maximum number of loci. For these GWAS (height, BMI, WHRadjBMI, red blood cell count, white blood cell count, diastolic blood pressure, and systolic blood pressure), we used a cutoff of p<5×10^-8^.

As described, DEPICT’s default behavior is to prioritize based on correlation across the reconstituted gene sets. The implicit assumption in this method is that the genes most likely to be truly causal are the ones with the most similar profile across these gene sets. As a first question for Benchmarker, we asked: how would correlating across tissue expression (rather than gene set membership) fare in comparison? To answer this question, we applied DEPICT’s prioritization algorithm, exchanging the matrix of z-scores of reconstituted gene sets for a matrix of z-scores of expression data (gene expression for each gene across a range of tissues). We used two different expression data sources. The first was the matrix DEPICT typically uses to perform tissue enrichment analysis, which was derived from 37,427 publicly available human microarrays representing 209 different tissues (based on Medical Subject Heading annotations). We note that this is a subset of the microarray data from the Gene Expression Omnibus (GEO)^32^ used in the process of “reconstituting” the gene sets. The second source of expression data was a matrix based on RNA-seq data from the Genotype Tissue Expression project (GTEx, v6)^33^, including 53 human tissues with an average of 161.32 samples per tissue (processed as in Finucane et al. 2018^34^). For each tissue, we calculated the mean expression across all samples. To make the expression data as comparable as possible, we normalized the GTEx expression matrix in the same way as the GEO matrix^9^: z-score normalizing across all tissues, then across genes. For all DEPICT analyses, we considered the top-ranked 10% of genes “prioritized.” All analyses were done on a set of 16,876 genes that were 1) outside the major histocompatibility complex region (chromosome 6: 25-35 Mb in hg19 genome build) and 2) present in both the DEPICT and GTEx data.

We also evaluated an additional prioritization method, MAGMA (v1.06b)^35^. MAGMA works in two steps. First, a gene-based p-value is computed as the mean association of SNPs in the gene, corrected for LD. Then, competitive gene set and/or continuous covariate p-values are calculated based on the association of the gene-based p-values with the category of interest. We ran MAGMA with default parameters. For gene set enrichment analysis, we treated the reconstituted gene sets as a continuous covariate and calculated one-tailed p-values (alternative hypothesis = genes with high gene set membership z-scores have a stronger trait association than those with low gene set membership z-scores).

Comparing DEPICT to MAGMA presents an immediate challenge: MAGMA does not explicitly prioritize genes based on the gene set enrichment analysis, as DEPICT does. Therefore, we needed to establish a framework for deriving gene prioritization from gene set enrichment analysis results (Supplemental Figure 1). Specifically, we first generated gene-based p-values from MAGMA. Then, we removed all genes from each chromosome in turn and applied the gene set enrichment analysis function to the remaining genes. From the gene set enrichment analysis results, we reasoned that we could prioritize genes that were members of the most highly enriched gene sets. However, the reconstituted gene sets used for the gene set enrichment do not have “members” per se; rather, they have z-scores for gene set membership prediction. To address this issue, we created several binarized forms of the gene sets in which we used z-score cutoffs (z > 1.96, 2.58, or 3.29, which correspond to 2-tailed p-values of .05, .01, and .001, respectively) or rankings (top 50, 100, or 200 genes per gene set) to define the gene set “members” (Supplemental Figure 2). (For the z > 3.29 condition, we removed 14 gene sets that contained fewer than 10 genes.) Therefore, for each withhold-one-chromosome gene set enrichment analysis result, we 1) ranked the gene sets and 2) for each gene set, annotated each member gene on the withheld chromosome as “prioritized” until we reached 10% of genes on the withheld chromosome (where gene set members were based on one of the six versions of the binarized gene sets). For the purposes of comparison, we applied DEPICT in exactly the same way, performing the prioritization based directly on the leave-one-chromosome-out gene set enrichment results and the binarized gene sets. We note that this basic strategy is itself a useful technique for converting the results of any gene set enrichment analysis to gene prioritization; this may be helpful in generalizing the types of data that can be used with Benchmarker.

## Results

The Benchmarker framework is outlined in Figure 1. First, we remove one chromosome from a set of GWAS summary statistics. Then, we apply a gene prioritization method of interest to this partial GWAS; this produces prioritization p-values for each gene in the genome, including on the withheld chromosome. We rank the genes on the withheld chromosome by prioritization p-value and take the top 10% as “prioritized.” Then, we annotate all SNPs within +/- 50 kb of these genes as prioritized SNPs. We repeat this process for each chromosome, successively withholding each chromosome and annotating prioritized SNPs. We then combine all the prioritized SNPs for each chromosome into a single “prioritized” annotation. Finally, we apply stratified LD score regression, which produces an estimate of the average per-SNP heritability (τ) of the prioritized annotation. For each of our applications, we tested GWAS of 20 different traits; to improve comparability across traits, we normalize our estimates of τ by the average genome-wide per-SNP heritability of each trait. We note that this method can easily be used for variant prioritization strategies in addition to gene prioritization strategies, with the only difference being that prioritized variants on the withheld chromosome can be directly annotated for stratified LD score regression without the step of mapping variants to genes.

We first assessed whether the type 1 error rate of our method was well-controlled by conducting 1000 null simulations using randomly prioritized SNPs. For each simulation, we randomly selected 10% of the SNPs that were within +/- 50 kb of any gene on each chromosome. These annotations were then used as input to stratified LD score regression. We tested each simulation against summary statistics from 10 different well-powered GWAS (for a total of 10,000 runs: 1000 null simulations each using 10 GWAS), including BMI, diastolic blood pressure, age of menopause, height, schizophrenia, years of education, age of menarche, IBD, total cholesterol, and allergy/eczema. The coefficient z-scores were well-controlled, with a type 1 error rate of 0.048 at p = .05 (Supplemental Figure 3a), indicating that stratified LD score regression is well-calibrated and correctly determines that a group of randomly selected SNPs does not significantly explain any heritability in any of the tested GWAS (i.e. in actual GWAS results). To more explicitly mimic our experimental setup, we also tested type 1 error by randomly sampling 10% of genes rather than variants (again, 1000 null simulations each using 10 GWAS for a total of 10,000 runs). Then, we annotated all SNPs within +/- 50 kb of these genes as “prioritized.” We observed that in this case, the estimates were actually slightly overconservative, with coefficient z-scores having a slight negative bias (type 1 error rate at 0.05 = 0.0593) (Supplemental Figure 3b). We speculate that this may result from the fact that SNPs within +/- 50 kb of genes have a nonrandom minor allele frequency distribution, which could affect the LD score regression estimates. Given that the bias was small and slightly conservative (in the negative direction for heritability explained), we proceeded without additional correction.

Next, we applied Benchmarker to compare three different methods of running DEPICT’s prioritization algorithm: a) prioritizing based on shared patterns of gene set membership (DEPICT’s standard approach, using a gene x gene set matrix of z-scores) and b) prioritizing based on shared patterns of tissue expression, using either of two different gene x tissue expression matrices of z-scores (one microarray-based from GEO^32^ and one RNA-seq based from GTEx^33^; see Methods). We refer to these approaches as DEPICT-gene-sets, DEPICT-GEO, and DEPICT-GTEx, respectively. We observed that the prioritized variants had coefficients significantly greater than 0 for nearly all traits and all three input matrices, indicating that prioritization by DEPICT using any of these data sources captures meaningful additional heritability beyond the effects of the baseline model (Supplemental Figure 4; Supplemental Table 2). However, within each trait, there were few clearly significant differences between the coefficients and no obvious overall trend for the three methods, indicating that the three methods were all performing similarly.

All three of these methods use gene expression information as a data source, either explicitly (for the two gene x tissue expression matrices) or implicitly (for the gene x gene set matrix, where z-scores for gene set membership are derived from gene expression data). We therefore considered whether the similarity in performance across methods might arise because the three methods prioritize similar genes. However, when we compared the overlap of prioritized genes, we found that in fact the approaches prioritized relatively distinct groups of genes. Specifically, the average number of genes prioritized by all three methods for a given trait was 374.1; in contrast, the average number of genes prioritized by the DEPICT-gene-sets only, DEPICT-GEO only, and DEPICT-GTEx only were 796.7, 732.4, and 757.7, respectively (Figure 2). Therefore, the use of three different data sources produced three substantially different groups of genes that each have similar contributions to heritability, suggesting that each set of prioritized genes contains different but equally useful information.

**Figure 2:**
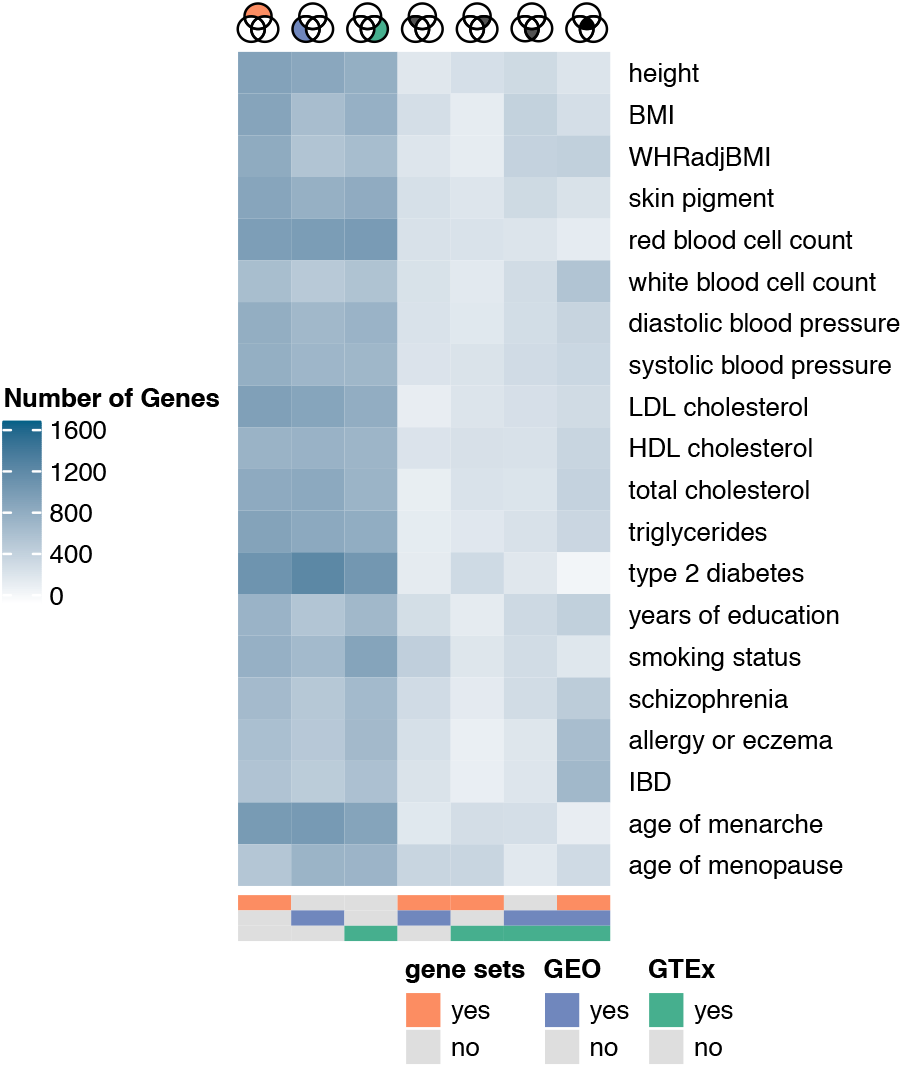
Overlap in prioritized genes for DEPICT-gene-sets, DEPICT-GEO, and DEPICT-GTEx. Columns of the heat map represent all possible categories of overlap, also illustrated by the Venn diagrams on top and the annotation bar below (e.g. prioritized by DEPICT-gene-sets only, prioritized by DEPICT-gene-sets and DEPICT-GEO, prioritized by all three methods, etc.). Darker blues indicate more genes in the category.

Because of this partial overlap in prioritized genes across the three methods, we tested whether combining the results from these methods might produce better prioritizations than any one method alone. Specifically, we created a new annotation consisting of genes prioritized by at least two of the three input matrices, which we refer to as the “intersect” set (average number of genes = 1203.05; average proportion of SNPs = 0.072) (Supplemental Figure 5, 6). We also created a separate annotation consisting of the remaining prioritized genes, i.e. genes prioritized by only one of those three methods; for simplicity, we refer to this group as the “outersect” set (average number of genes = 2286.80; average proportion of SNPs = 0.120). We first tested the performance of the “intersect” and “outersect” annotations in separate LD score regression models (Supplemental Figure 4; Supplemental Table 2) and found that across most traits, the intersect performed better than the outersect, in some cases significantly. To more directly compare these annotations, we performed a conditional analysis in which the annotations were modeled jointly (i.e. in the same LD score regression model) (Figure 3; Supplemental Table 3). We observed a clear difference between the two annotations: across most traits, the intersect group of genes substantially outperformed the outersect (random-effects metaanalysis p for 16 traits = 1.39 x 10^-4^; we excluded four phenotypically overlapping traits, see Methods). These results indicate that, while all three methods performed similarly, the majority of the heritability explained could actually be localized to SNPs near or within the genes prioritized by more than one method; moreover, in most cases, the intersect group of genes was smaller than the outersect group, implying that the intersect identifies a smaller group of genes with a higher enrichment for heritability.

**Figure 3:**
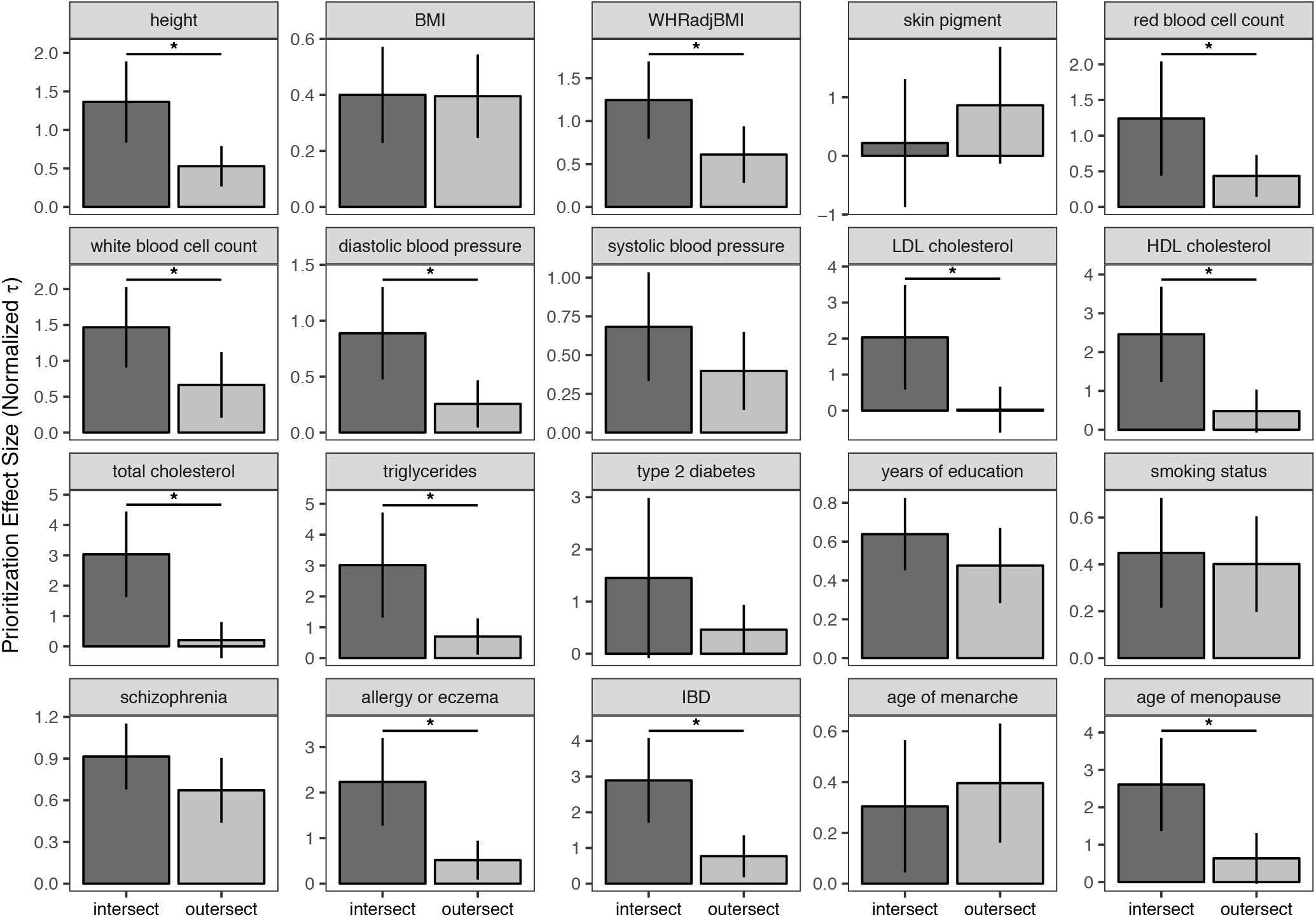
Effect sizes (normalized τ) for the joint LD score regression model comparing “intersect” and “outersect” genes for 20 different GWAS. Here, the “intersect” represents genes prioritized by at least two of 1) DEPICT-gene-sets, 2) DEPICT-GEO, and 3) DEPICT-GTEx. The “outersect” represents genes prioritized by only one of those three methods. Asterisks mark comparisons for which the difference between the intersect and outersect achieved nominal significance (p < 0.05). Error bars represent 95% confidence intervals. Note that y-axis scales differ for each trait.

We also noted that the intersect outperformed the outersect most strongly for the lipid traits (LDL and HDL cholesterol, total cholesterol, and triglycerides), immune traits (allergy/eczema and IBD), and height. In contrast, we observed that the most brain-related traits we tested (BMI, years of education, smoking status, schizophrenia, and age of menarche) failed to show a nominally significant difference between the intersect and outersect (p > 0.05). This suggests that brain-related traits may not benefit as much as other traits from combining information across these particular data sources (i.e. the reconstituted gene sets and tissue expression matrices).

We next wanted to compare DEPICT with another popular gene set enrichment analysis algorithm, MAGMA^35^. However, MAGMA does not perform gene prioritization based on its gene set enrichment, so we needed a way to convert prioritized gene sets to prioritized genes. We accomplished this by (1) ranking the prioritized gene sets, (2) using binarized versions of the reconstituted gene sets to assign genes to gene sets, and (3) prioritizing genes in the most enriched gene sets on the withheld chromosome (Supplemental Figure 1). For step 2, we created six different binarized versions of the reconstituted gene sets, three based on z-score (z > 1.96, 2.58, or 3.29) and three based on ranking (top 50, 100, or 200 genes per gene set). For the purposes of comparison, we used the same strategy for DEPICT (i.e. using the binarized gene sets to identify and prioritize genes on the withheld chromosome within enriched gene sets). This basic approach can be used for any method that prioritizes genomic features (e.g. gene sets, tissue expression, epigenomic annotations) but does not necessarily explicitly prioritize genes or variants; it also illustrates that the Benchmarker strategy can be used to evaluate a wide variety of algorithm types.

DEPICT and MAGMA performed similarly: we observed no strongly significant differences between the two methods for each trait (Supplemental Figure 7; Supplemental Table 4). However, as we observed for the comparison across different data sources, we again noted that DEPICT and MAGMA prioritized fairly different groups of genes (average number of genes prioritized by both methods = 931.18, average number of genes prioritized by one method only = 757.82) (Figure 4, Supplemental Figures 8-10). Using the same logic as before, we again asked whether the genes found by both methods (the “intersect”) outperformed the genes found by either method individually (the “outersect”). (Note that the outersect includes the union of genes prioritized by only DEPICT and only MAGMA, so it is on average slightly larger than the intersect [Supplemental Figure 10].) Interestingly, we observed an even stronger trend toward the overlapping set outperforming the individual sets than for the analysis of different data sources, and for several traits this difference was nominally significant (Supplemental Figure 7; Supplemental Table 4). When we modeled the intersect and outersect jointly, this difference became even more apparent, with the intersecting group of genes outperforming the outersect for nearly every trait (Table 1, Figure 5, Supplemental Table 5). The differences were particularly pronounced for immune traits (IBD and allergy/eczema) and total cholesterol (which, interestingly, also were some of the best-performing traits in the previous analysis). For these traits, not only did the intersect significantly outperform the outersect for every gene set binarization, but outersect per-SNP heritability often did not significantly differ from zero. This implies that the vast majority of the heritability originally explained by both sets of prioritized genes (i.e. the union of DEPICT and MAGMA) was captured by the intersect genes only, with almost none remaining in the outersect.

**Figure 4:**
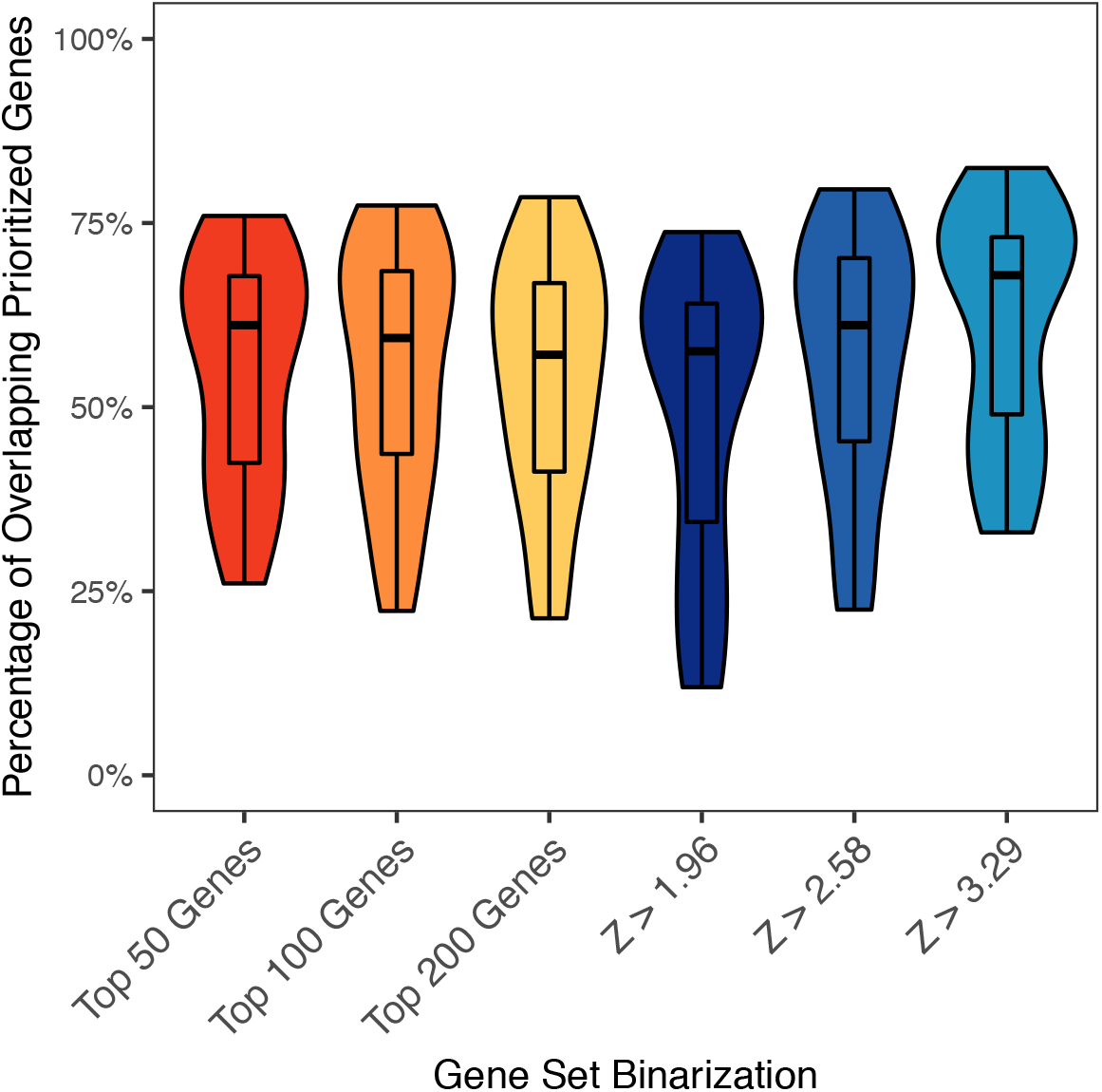
Overlap in prioritized genes for DEPICT versus MAGMA. For each version of the binarized gene sets (x-axis), the distribution of the percentage of overlapping genes across all 20 traits is represented as a violin plot overlaid with a box plot.

**Table 1.** Summary of results for comparison of “intersect” and “outersect” genes for DEPICT and MAGMA. Each row includes data based on one of the six gene set binarizations. The columns represent the overall p-value for comparing τ_intersect_ and τ_outersect_ and results from the height OMIM enrichment analysis (odds ratio and p-value from Fisher’s exact test). OMIM = Online Mendelian Inheritance in Man.

| Gene Set Binarization | Meta-analysis p-value for $\tau_{\text{intersect}}$ versus $\tau_{\text{outersect}}$ | Height OMIM enrichment odds ratio (intersect versus outersect) | Height OMIM enrichment p-value (intersect versus outersect) |
| --- | --- | --- | --- |
| Top 50 Genes | 8.547e-07 | 2.53 | 0.000154 |
| Top 100 Genes | 1.557e-05 | 2.421 | 0.000210 |
| Top 200 Genes | 1.084e-04 | 1.364 | 0.185 |
| Z > 1.96 | 2.537e-04 | 0.82 | 0.437 |
| Z > 2.58 | 8.489e-05 | 1.300 | 0.243 |
| Z > 3.29 | 9.110e-06 | 1.870 | 0.000618 |

**Figure 5:**
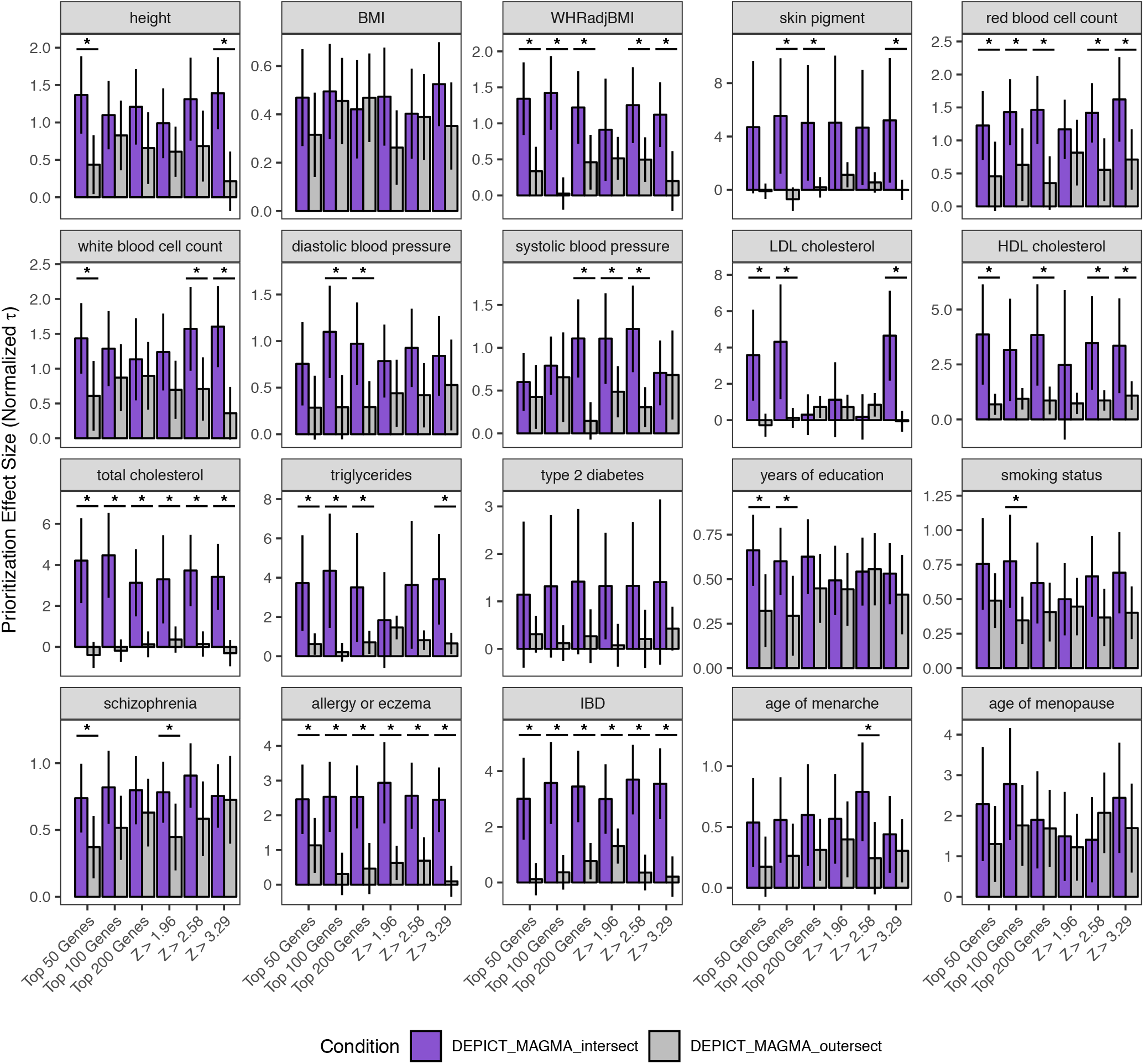
Effect sizes (normalized τ) for the joint LD score regression model comparing “intersect” and “outersect” genes for 20 different GWAS. Here, the “intersect” represents genes prioritized both DEPICT and MAGMA. The “outersect” represents genes prioritized by only DEPICT or only MAGMA. Asterisks mark comparisons for which the difference between the intersect and outersect achieved nominal significance (p < 0.05). Error bars represent 95% confidence intervals. Note that y-axis scales differ for each trait.

We also observed that the top 50 genes, top 100 genes, and Z > 3.29 conditions showed the strongest performance. These conditions also represent the gene sets with the smallest number of genes (Supplemental Figure 2), suggesting that for our choice of method, more stringent cutoffs for gene set membership produce more power. Given our prioritization schema (Supplemental Figure 1; see Methods), this is not surprising: when gene sets have a very large number of members, a few gene sets will dominate the gene prioritization results. In contrast, when smaller gene sets are used, more gene sets will end up contributing prioritized genes, likely beneficially increasing diversity.

As an additional validation for the intersection sets, we used a previously curated list of 277 genes from Online Mendelian Inheritance in Man known to cause disorders of skeletal growth^22^. We note that in general, we do not endorse gold standards, but we use them here as an orthogonal source of validation data for height, where there are a large number of well-established and well-validated Mendelian disease genes. We performed a Fisher’s exact test to determine whether the intersect genes for height were enriched for these genes relative to the outersect genes. We found that in the gene set versus tissue analysis, the intersect was indeed enriched for OMIM genes (odds ratio = 2.004, p = 4.72 x 10^-4^). Similarly, in the three versions of the binarized gene sets with the most significant meta-analysis p-value, we also observed an enrichment of OMIM genes (Table 1).

## Discussion

We have developed and implemented a leave-one-chromosome out approach for benchmarking gene prioritization algorithms. The use of explained heritability as a metric has several advantages over methods that have been used elsewhere^17,20^. Firstly, it does not rely on the idea of “true positive” or “gold standard” genes, which is inherent in commonly used area-under-the-curve benchmarks; true positives are only as good as the existing knowledge base on a given trait and are likely biased towards a subset of relevant biology. Secondly, heritability explained directly measures an actual quantity of interest, namely, how well the method can independently identify genes that colocalize with GWAS association signals. Our leave-one-chromosome-out strategy is also an obviously unbiased way of measuring method performance, as it enables the use of the GWAS data itself as its own control rather than external sources of data. We recommend that this idea should be used in benchmarking new and existing prioritization methods; that is, testing should be done on genes or variants prioritized on a chromosome (or a set of chromosomes) that has been withheld from the input data. We also recommend leaving out at least an entire chromosome to avoid overfitting from correlation of association signals from neighboring genes. In this way, it is possible to both have the advantages of cross-validation without the disadvantages of relying on gold standards.

In two different sets of comparisons, one using different data sources, and one using different prioritization algorithms, we showed that the performance of different approaches is relatively similar, but that selecting genes prioritized by multiple approaches outperforms the use of genes prioritized by any individual approach. Supporting this observation, Mendelian genes for skeletal growth disorders are more enriched in “intersect” than “outersect” genes for height. This finding has important implications for translating genetic associations into biological insights, as it empirically demonstrates that combining prioritization approaches is superior to relying on the somewhat arbitrary choice of a single approach. We also observed that intersect performance was particularly strong for immune and lipid traits. In contrast, brain-related traits generally showed fewer significant differences between intersect and outersect gene performance. One possible explanation is the heterogeneity of the brain, which has extremely high regional and cell-type specificity. Gene prioritization for these traits might therefore improve with a different approach, such as restricting tissue expression data to brain regions only, or, conversely, including only a single representative brain region in the tissue expression analysis rather than many. Another possible reason for this finding is the importance of brain-region-specific isoform expression, which is not accounted for in our general tissue expression data.

We also note that, because Benchmarker relies on a cross-validation strategy, it can fairly be used to determine the best prioritization method for any given trait. This strategy provides a route to new best practices for gene prioritization for the field: by benchmarking multiple approaches (and particularly the intersection of multiple approaches), it should be possible to objectively improve the gene prioritization of any given trait. For example, we observed that age of menarche was one of the few traits for which combining information across gene-set- and tissue matrix-based prioritization did not improve per-SNP heritability; one possible explanation based on our original analysis is that the GTEx matrix alone did not significantly contribute to heritability, so using information from that analysis may have actually worsened the signal-to-noise ratio. Such observations have the potential to inform specific decisions for performing gene prioritization for individual traits. In addition, with our analysis of MAGMA, we have shown that this benchmarking strategy can be extended to methods that evaluate enrichment of genomic features (e.g. pathways or tissue expression) but do not explicitly assign prioritization p-values to genes. This high generalizability will allow for the comparison of a wide variety of approaches for different traits.

An important caveat to our results is that LD score regression may have insufficient power to identify small differences in explained heritability using different approaches to gene prioritization, and that other metrics (such as the genomic inflation factor for prioritized variants) may be more sensitive. We note, however, that LD score regression is a widely-used method in the field and has found important and significant differences in explained heritability by a variety of other types of annotation, such as cell-type-specific expression^34^ and epigenomic marks^21^. We believe that any small losses in power from our choice of method are outweighed by the benefits of the output (i.e. per-SNP heritability) being directly interpretable and extremely meaningful. Furthermore, improvements to the annotations we have used here will provide boosts in power (as well as answers to other questions about how effective such “improvements” actually are). For example, using a 50-kb window around prioritized genes, as we have done here, means that a large amount of noise is included in our annotations. A major improvement would therefore be assignment of noncoding SNPs to the genes they regulate based on expression, epigenetic, and/or chromatin conformation data.

In conclusion, we have developed a powerful and well-controlled approach for benchmarking gene prioritization strategies that relies solely on GWAS data and does not require any assumptions about the “correct” biology. Our method shows that combining prioritization strategies produces smaller groups of genes that are more enriched for heritability than any strategy alone and suggests a strong recommendation that follow-up studies be focused on genes prioritized using multiple approaches. Future methods will benefit from incorporating different statistical approaches and data sources; even apparently similar data (such as two different sources of tissue expression) can provide different and complementary information. We believe that the overall cross-validation approach described and implemented here provides a better “gold standard” for benchmarking existing and future methods for gene and variant prioritization. Finally, our method can be used to determine the best algorithm and data set for any particular trait of interest.

## Description of Supplemental Data

Supplemental Data include ten figures and five tables.

## Declaration of Interests

J.N.H. is on the scientific advisory board of Camp4 Therapeutics.

## Acknowledgements

We gratefully acknowledge Hilary Finucane, Steven Gazal, and Jack Kosmicki for helpful discussion. This work was supported by NHGRI F31HG009850 (R.S.F.), NIDDK NIH R01DK075787 (J.N.H.), the Lundbeck Foundation R190-2014-3904 and the Novo Nordisk Foundation (T.H.P.), NHGRI NIH T32 HG002295 (T.A.), and NIAMS NIH 1R01AR063759-01A1 (S.R.).

## Web Resources

Benchmarker: https://github.com/RebeccaFine/benchmarker

LDSC software: https://github.com/bulik/ldsc (version 1.0)

LD scores and other files for LDSC: https://data.broadinstitute.org/alkesgroup/LDSCORE/

DEPICT software: https://data.broadinstitute.org/mpg/depict/ (release 194)

MAGMA software: https://ctg.cncr.nl/software/magma (version 1.06b)

GTEx: http://www.gtexportal.org

